# Presence of *Toxoplasma gondii* tissue cysts in human semen

**DOI:** 10.1101/2021.10.27.466215

**Authors:** Wen Han Tong, Jana Hlaváčová, Samira Abdulai-Saiku, Šárka Kaňková, Jaroslav Flegr, Ajai Vyas

**Affiliations:** School of Biological Sciences, Nanyang Technological University, 60 Nanyang Drive, Singapore – 637551; Department of Philosophy and History of Science, Faculty of Science, Charles University, Viničná 7, CZ – 128 44 Prague, Czech Republic

**Keywords:** *Toxoplasma gondii*, Sexual transmission, Toxoplasmosis, Venereal disease, Blood-testes barrier, Coccidian parasite

## Abstract

*Toxoplasma gondii* is a widely prevalent protozoan parasite in human populations. This parasite is thought to be primarily transmitted through undercooked meat and contamination by cat feces. Here, we demonstrate that *Toxoplasma gondii* cysts can be found within human semen, thus suggesting a potential for sexual transmission. We visualized *Toxoplasma gondii* cysts in ejaculates of immune-competent and latently infected human volunteers. We confirmed the encystment by probing transcription of a bradyzoite-specific gene in these structures. These observations extend previous observations of the parasite in semen of several non-human host species, including rats, dogs, and sheep. *Toxoplasma gondii* infection is a clinically significant infection, in view of its high prevalence, its purported role in neuropsychiatric disorders such as schizophrenia, as well as in the more serious form of congenital toxoplasmosis. Our demonstration of intact *Toxoplasma gondii* cysts in the ejaculate supports the possibility of sexual transmission of the parasite and provides an impetus for further investigations.

**Importance:** *Toxoplasma gondii* is recognized as one of the five infections that can harm fetal health. There is also emerging evidence that toxoplasmosis is related to the development of neuropsychiatric disorders. In this context, the current report includes visual evidence of *Toxoplasma gondii* cyst presence in the semen of human males. It is thus plausible that this clinically significant infection can be sexually transmitted. The existence of sexual transmission in a population, and its relative importance vis-à-vis other transmission routes, remains undetermined. These possibilities are of acute relevance to human health in the context of sexual health and pregnancy management.

## Introduction

*Toxoplasma gondii* (thereafter *Toxoplasma*) is a protozoan and intracellular parasite with a two-stage life (1). Its range of intermediate hosts includes a wide variety of homeothermic animals, including livestock, humans, and human commensals. Sexual reproduction or gametogenesis is confined to the gut of members of the cat family Felidae (2). *Toxoplasma* is one of the most prevalent human parasitic infections, with epidemiological studies suggesting a prevalence rate of ∼30-40% worldwide (3). Prevalence is estimated to be ∼11% in the USA and 17% in Singapore, with some geographical locations reporting a prevalence rate as high as 70% (3-5). *Toxoplasma* differentiates into fast-dividing tachyzoites during the acute phase of the infection, disseminating in a wide range of tissue types. As the host body mounts immune response and inflammation, *Toxoplasma* migrates to immune-privileged sites like the brain and eyes. Here it differentiates into bradyzoites housed within tissue cysts, characterized by slow division and quiescent metabolism (1). These tissue cysts go through periodic recrudescence and cycles of immune response and re-encystment. The serological presence of anti-*Toxoplasma* IgG antibodies is often used as a proxy for the presence of latent *Toxoplasma* infection. Immune-competent humans typically do not display overt clinical symptoms (6). Therefore, toxoplasmosis is commonly thought of as a latent and clinically asymptomatic infection with lifelong persistence. However, this perspective is undergoing revision at present. Several reports of retrospective and prospective studies associate *Toxoplasma* infection status with an increased risk of developing neuropsychiatric disorders including bipolar disorder, addiction, obsessive compulsive disorder and schizophrenia (7-13).

Humans are primarily infected through exposure to cat feces or by consuming undercooked meat containing parasitic cysts (14). Vertical transmission from acutely infected mothers to the developing fetus and organ transplants are other possible modes of *Toxoplasma* transmission in humans (15). Interestingly, there is growing evidence to non-human hosts that *Toxoplasma* can also be sexually transmissible. For example, *Toxoplasma* breaches the blood-testes barrier in rats and encysts within epididymis during latent infection (16, 17). *Toxoplasma* in this host species transfers to the females through ejaculate and establishes chronic infection in both females and resultant offspring. Aside from rats, viable parasites have been isolated from the semen of asymptomatic seropositive dogs (18) and goats (19). Subsequently, females artificially inseminated with this infected semen test positive for anti-*Toxoplasma* antibodies. Seroconversion is also achieved when female sheep are inseminated with *Toxoplasma*-spiked semen (20). These strands of evidence in non-human species suggest that *Toxoplasma* can anterogradely cross the blood-testes barrier, an uncommon ability amongst pathogens due to the highly impervious nature of this tissue barrier.

Recently, epidemiological and therefore indirect evidence of male-to-female sexual transmission in humans has been published. A study of sexual partners showed that seropositivity in men increases the risk of infection in women (21). There are strong indices that oral sex could be an important source of infection in heterosexual women and homosexual men (22).

Against this backdrop, we examined the possibility that live *Toxoplasma* parasites can be ejaculated along with semen in immunocompetent human subjects by examining the semen of seropositive immunocompetent men for the presence of the tissue cysts.

## Materials and Methods

### Sample collection

Human volunteers were recruited from the Center for Assisted Reproduction, Department of Obstetrics and Gynaecology of the First Faculty of Medicine of Charles University and General University Hospital in Prague, Czech Republic. The goal and experimental procedures of this study were explained to all participants. Volunteers signed informed consent before the participation. The mean age of the participants was 36 years (SD = 5.14; range: 26 – 54 years). Ethical approvals were obtained from institutional review boards in General University Hospital in Prague (#384/16; 92/17) and Faculty of Science, Charles University (#2015/29).

Venous blood and semen samples were collected under aseptic conditions. Serum was separated from whole blood via centrifugation (2500 rpm for 10 minutes) and stored at -20°C before analysis. The full set of samples comprised 723 men, of which 172 were found to contain anti-*Toxoplasma* IgG antibodies (23.8%). Several pairs of semen and matched serum samples were shipped to Singapore on dry ice after obliterating volunteer identity. These shipments contained a subset of the collected samples and were artificially enriched for seropositive participants. The serological status was anonymized upon shipping. All procedures conducted in Nanyang Technological University were reviewed and approved by the institutional review board (#IRB-2017-11-021).

### Serology

Serum was examined for the presence of anti-*Toxoplasma* IgG antibodies using commercial ELISA kits (Abcam). In line with the manufacturer’s recommendation, the amount of anti-*Toxoplasma* IgG antibody above ≥ 35 U/mL was considered seropositive, and values < 30 U/mL were categorized as seronegative. Values between 30 to 35 U/mL were classified as equivocal and inconclusive.

### Giemsa stain

5μl of semen sample was pipetted onto a Superfrosted glass slide (Fisher Scientific) after thawing. The semen sample was smeared across the glass slide and allowed to air-dry completely for an hour. Semen smear slides were stored at 4°C prior to usage.

Giemsa stain was carried out as previously described (23). In brief, semen-smeared slides were fixed for 10 minutes in 100% methanol at room temperature and then completely air-dried. The slides were then stained by Giemsa stain (Sigma-Aldrich) for 45 minutes at room temperature. The working Giemsa stain was prepared at a 1:5 dilution in phosphate-buffered saline and passed through a 0.45μm syringe filter. After staining, the slides were rinsed with deionized water and air-dried completely. Slides were coverslipped with a mountant and visualized in a microscope (Zeiss Live Cell observer light microscope, 40X objective lens with a 1.2X digital magnification; total magnification = 400X).

### Fluorescence immunohistochemistry

Fluorescence immunohistochemistry was carried out as previously described (24). The human semen smear slides were fixed in 4% paraformaldehyde (wt/vol, in buffered saline) for 30 minutes at room temperature. Slides were then permeabilized in buffered saline containing 0.25% (v/v) Triton X-100 for 20 minutes. Slides were blocked with bovine serum albumin for 1 hour at room temperature (Sigma-Aldrich, 1% v/v).

1X PBS containing 1% (vol/vol) Bovine Serum Albumin (Sigma-Aldrich, #A7906) for 1 hour at room temperature. Smear slides were then incubated with rhodamine-labeled *Dolichos biflorus* agglutinin (Vector Laboratories, dilution 1:100), a lectin that binds to carbohydrates found in the *Toxoplasma* tissue cyst wall. Slides were then repeatedly washed in buffered saline and air-dried. Slides were subsequently coverslipped with a mountant containing DAPI to stain nuclear boundaries (Life Technologies). Tissue cysts were visualized with a laser scanning confocal microscope (Carl Zeiss LSM 710), using a 40X objective lens with a 1.2X digital magnification (total magnification = 400X).

### In situ hybridization

In-situ hybridization was performed on semen smears to visualize messenger RNA known to be specifically transcribed from the *Toxoplasma* genome in bradyzoites within tissue cysts (BAG1 mRNA sequence; GenBank: X82213.1). RNAScope probes against BAG1 mRNA were commercially obtained (ACDbio) using proprietary techniques. Glass-mounted semen smears were stained in accordance with supplier instruction, made available with the probes. A minus-probe negative control was included. Slides were scanned using the Zeiss Live Cell observer light microscope using a 40X objective lens with a 1.2X digital magnification (total magnification = 400X).

## Results

### Histological visualization confirmed the presence of *Toxoplasma* cysts in the semen

Initially, Giemsa stain was used to visualize *Toxoplasma* cysts in semen smears. *Toxoplasma* cysts can be easily identified in this staining preparation, characterized by a heavily stained interior containing bradyzoites along with a lightly stained cyst wall (Figure 1A and 1B) (23). A subset of semen from fifty volunteers, previously determined to be seropositive for IgG antibodies, was used for this experiment.

**Figure 1.**
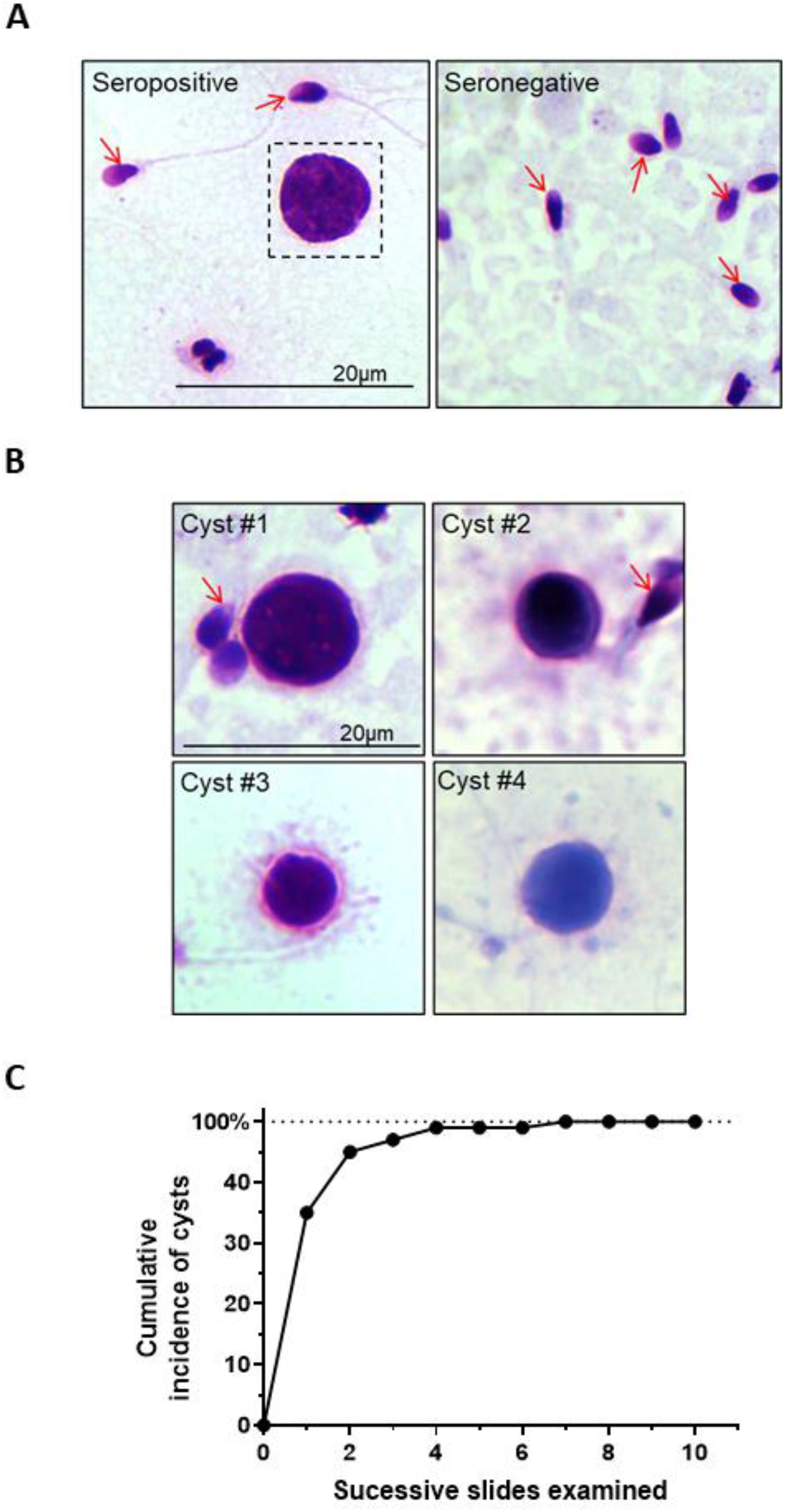
**(A)** Representative brightfield image depicting Giemsa-stained preparation of a *Toxoplasma* cyst within semen from a seropositive human male and a negative control. Sperm cells are denoted by red arrows. Scale bar = 20μm. **(B)** Magnified view of *Toxoplasma* cyst. **(C)** Cumulative distributions of Toxoplasma-seropositive volunteers showing microscopic evidence of Toxoplasma cysts. All 50 samples showed the presence of Toxoplasma cysts as an increasing number of semen slides was examined, reaching an asymptote at 100% concordance between serological presence of anti-Toxoplasma IgG antibodies and physical presence of Toxoplasma cysts in the semen.

Ten glass slides were prepared by smearing a total of 50 μl of the sample. The presence of *Toxoplasma* was apparent in 35 out of 50 cases when the first slide from each individual was stained and visualized containing five μl of semen. We then stained an additional slide for 15 cases that did not show the presence of the parasite in the first instance, noting the presence of *Toxoplasma* in further ten cases. Upon continuing this successive examination, all 50 seropositive cases contained *Toxoplasma* cysts in ≥ 1 slide (Figure 1C).

We subsequently sought to confirm the identity of *Toxoplasma* cysts by staining the cyst wall with a lectin with a known affinity for wall carbohydrates (24). A rhodamine-labeled *Dolichos biflorus* agglutinin was used for this purpose. Differential interference contrast showed evidence of fluorescent cyst wall encompassing spherical cyst-like structures in the semen smears (Figure 2). We further probed a bradyzoite-specific mRNA to confirm that the intra-cyst structures indeed constituted *Toxoplasma* bradyzoites. In-site hybridization confirmed the presence of BAG-1 mRNA inside the tissue cysts, specifically demonstrating the transcription event specific to *Toxoplasma* encystment (Figure 3).

**Figure 2.**
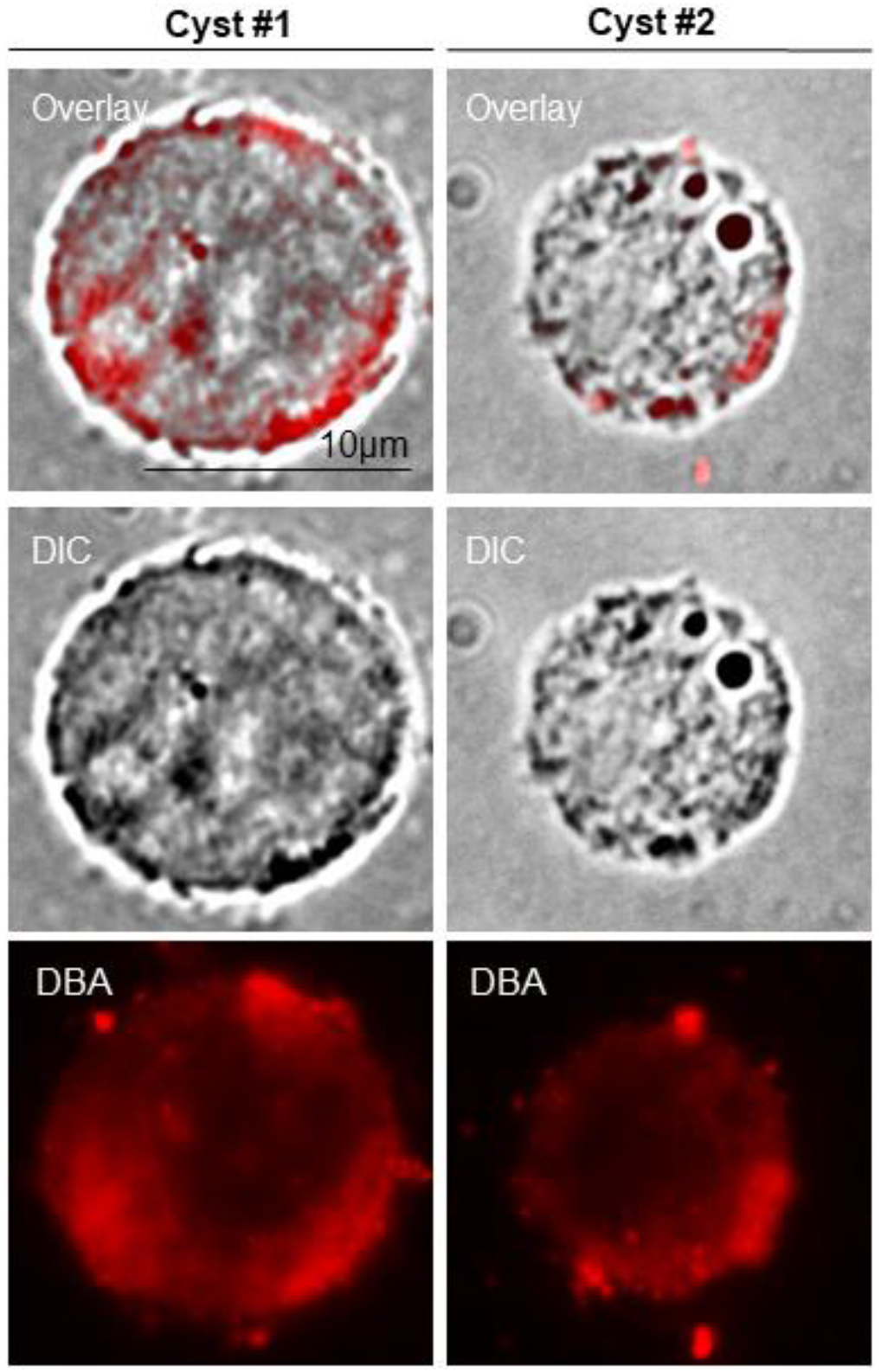
Representative confocal images depicting *Toxoplasma* cyst wall (red), stained with rhodamine-labeled *Dolichos biflorus* lectin, superimposed on a brightfield differential interference contrast image of the cyst. Scale bar = 10μm.

**Figure 3.**
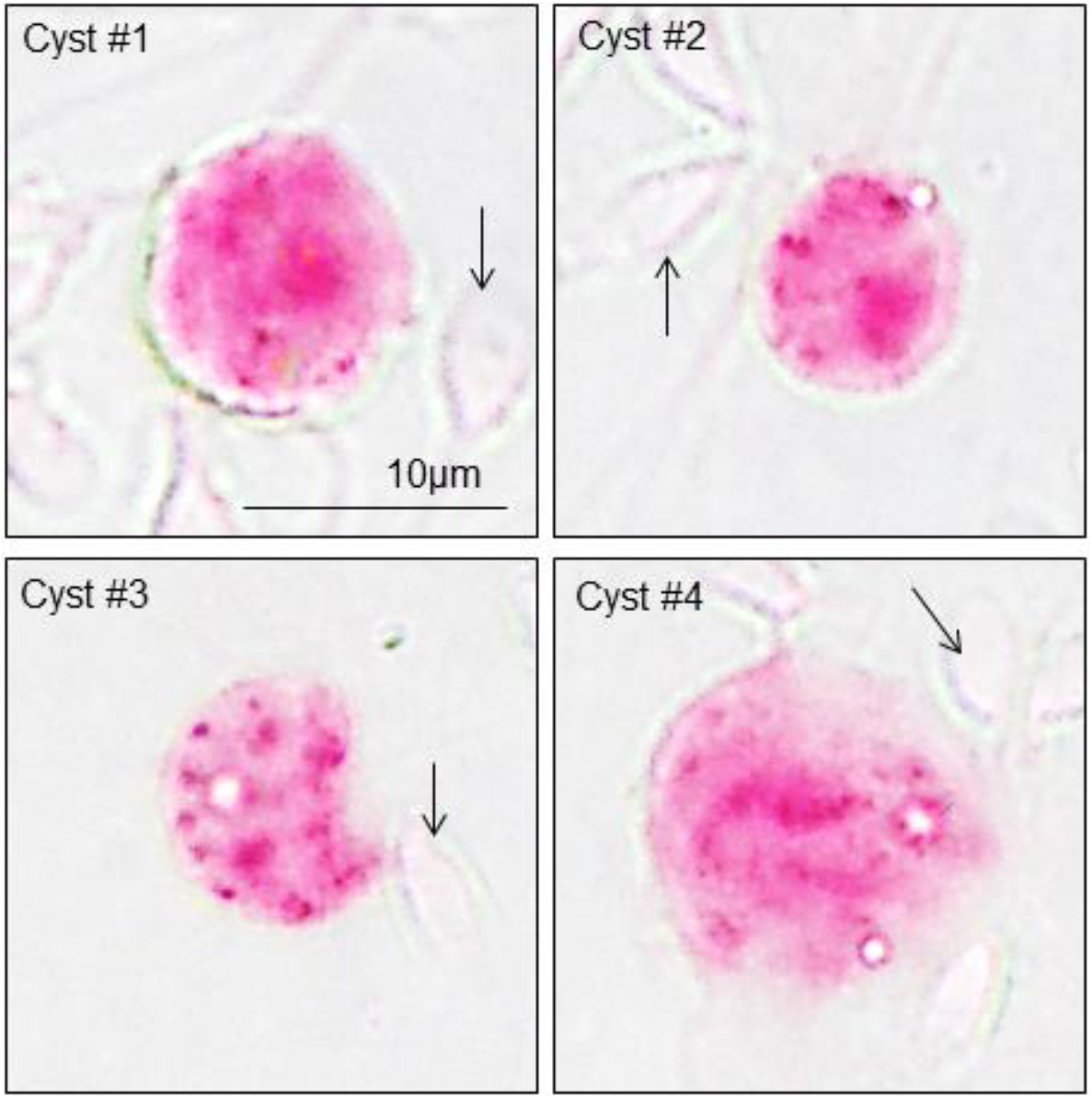
Transcription of bradyzoite-specific BAG-1 mRNA within tissue cysts visualized by RNAScope technology. *Toxoplasma* bradyzoites are stained in red chromogenic dye. Bradyzoites are contained and surrounded by the relatively thin cyst wall. Sperm cells are denoted by black arrows. Scale bar = 10μm.

## Discussion

In this report, for the first time, we demonstrate the presence of *Toxoplasma* tissue cysts within human semen. These observations support the notion that *Toxoplasma* is a sexually transmissible infection in humans, congruent to prior observations made in hon-human intermediate hosts.

Although infections in the human population are mainly asymptomatic (but see (25, 26)), toxoplasmosis remains a public health problem for some populations. Symptomatic congenital infection by *Toxoplasma*, for example, presents with a prevalence rate of 0.7 cases per 1000 live births in the USA (27). Moreover, the infection during the first trimester of pregnancy in expectant mothers who have not been previously infected by *Toxoplasma* carries a substantial risk of miscarriages or stillbirth (28). The existence of the venereal route of the *Toxoplasma* transmission could potentially explain the poor efficiency of the educational campaigns that strive to reduce congenital toxoplasmosis risk. A large fraction of *T. gondii* infections in pregnant women cannot be attributed to the known risk factors (29, 30), and sexual transmission could explain part of these infections. The potential of sexual transmission, thus, can signify particular risk in terms of pregnancy outcome and is also of consequence to the operation of sperm banks.

Our observations show the presence of tissue cysts in the semen but do not prove if these cysts can successfully initiate and establish infection in sexual partners post-coitus. Earlier non-human studies have demonstrated that ejaculated *Toxoplasma* cysts can result in seroconversion in females as a direct result of intercourse (16, 31, 32). Such clear data remainsabsent in the case of humans, although several strands of evidence support that possibility. A recent study using a similar cohort of clients from the Center of Assisted Reproduction, for example, showed that *Toxoplasma* infection was 1.4 times more prevalent in women partners of the infected men at a prevalence rate of 25.6%, compared to partners of uninfected men at a rate of 18.2% women (21). Concordance of serostatus between sexual partners cannot, by itself, provide strong support to the possibility of sexual transmission. Partners often share living environments and other risk factors like exposure to cats, soil, or cooking practices. Nonetheless, the increased risk of the infection due to living with infected partners appears specific to women and not men. This sexual dichotomy supports the idea of sexual transmission through ejaculate (21). Moreover, there is a positive association between the prevalence of *Toxoplasma* infection and the prevalence of other sexually transmitted diseases in humans, which further consolidates the argument that *Toxoplasma* can be sexually transmitted (14). *Toxoplasma* cysts are resistant to, and possibly even need to be activated by, gastric juices, with consumption of undercooked meat being a primary risk factor for the infection. A recent study raises the possibility that oral sex can be a risk factor for *Toxoplasma* infection in heterosexual and homosexual partners (22). According to this, the seroprevalence of toxoplasmosis was increased in women and (non-significantly) also in men reporting to be engaged in oral sex (fellatio, not cunnilingus) [29].

The blood-testes barrier consists of Sertoli cells that are present at the exterior surfaces of seminiferous tubules and form inter-cellular tight junctions (33, 34). Two conducive but distinct microenvironments exist for the development of germ cells. The basal part of seminiferous tubules houses spermatogonial cell division, differentiation, and renewal of preleptotene (35).

The basal part is separated from the adluminal or apical region by the blood-testes barrier, creating a immune-privileged environment for spermiogenesis, meiosis, and spermiation (36, 37). The blood-testes barrier is one of the most impervious barriers within the mammalian body. Most infections found in human seminiferous tubules, epididymis, and testes represent an retrograde invasion from the penile canal (38, 39). Relevant examples of such retrograde infections include *Neisseria gonorrhoeae* or *Chlamydia trachomatis* (38, 39). Furthur example of an infection with retrograde ascent includes *Pseudomonas aeruginosa* (40). In contrast, the anterograde passage of pathogens from the testicular lumen side is much infrequent in immunocompetent males. *Toxoplasma* might provide an interesting exception to this rule. Congruent with the presence of the parasite within testes, *Toxoplasma* increases testosterone production in rats (41) and humans (42); and facilitates testosterone-dependent phenotypes in males (43-45).

Observations in this study lay the groundwork for future experiments to determine if the presence of the parasite in human semen can be generalized to other population cohorts. Further experiments are also warranted to examine the possibility and relative importance of sexual transmission in the epidemiology of *Toxoplasma* infection and the etiology of congenital toxoplasmosis.

## Declarations

## Acknowledgment

This work was financially supported by the Ministry of Education, Singapore (grant RG136/15) and Charles University Grant Agency (grant 262721).

## Statement of Ethics

All experimental procedures were reviewed and approved by Institutional review boards of University Hospital in Prague (# 384/16; 92/17), Faculty of Science at Charles University (#2015/29), and Nanyang Technological University (#IRB-2017-11-021).

## References

1. Black MW, Boothroyd JC. 2000. Lytic Cycle of Toxoplasma gondii. Microbiology and Molecular Biology Reviews 64:607–623.

2. Berdoy M, Webster JP, Macdonald DW. 2000. Fatal attraction in rats infected with Toxoplasma gondii. Proc Biol Sci 267:1591–4.

3. Jones JL, Kruszon-Moran D, Rivera HN, Price C, Wilkins PP. 2014. Toxoplasma gondii seroprevalence in the United States 2009-2010 and comparison with the past two decades. Am J Trop Med Hyg 90:1135–9.

4. Lim KC, Pillai R, Singh M. 1982. A study on the prevalence of antibodies to Toxoplasma gondii in Singapore. Southeast Asian J Trop Med Public Health 13:547–50.

5. Wong A, Tan KH, Tee CS, Yeo GS. 2000. Seroprevalence of cytomegalovirus, toxoplasma and parvovirus in pregnancy. Singapore Med J 41:151–5.

6. Flegr J. 2007. Effects of Toxoplasma on Human Behavior. Schizophrenia Bulletin 33:757–760.

7. Torrey EF, Bartko JJ, Lun ZR, Yolken RH. 2007. Antibodies to Toxoplasma gondii in patients with schizophrenia: a meta-analysis. Schizophr Bull 33:729–36.

8. Hinze-Selch D, Daubener W, Erdag S, Wilms S. 2010. The diagnosis of a personality disorder increases the likelihood for seropositivity to Toxoplasma gondii in psychiatric patients. Folia Parasitol (Praha) 57:129–35.

9. Prandota J. 2010. Autism spectrum disorders may be due to cerebral toxoplasmosis associated with chronic neuroinflammation causing persistent hypercytokinemia that resulted in an increased lipid peroxidation, oxidative stress, and depressed metabolism of endogenous and exogenous substances. Research in Autism Spectrum Disorders 4:119–155.

10. Miman O, Mutlu EA, Ozcan O, Atambay M, Karlidag R, Unal S. 2010. Is there any role of Toxoplasma gondii in the etiology of obsessive-compulsive disorder? Psychiatry Res 177:263–5.

11. Flegr J, Klose J, Novotna M, Berenreitterova M, Havlicek J. 2009. Increased incidence of traffic accidents in Toxoplasma-infected military drivers and protective effect RhD molecule revealed by a large-scale prospective cohort study. BMC Infect Dis 9:72.

12. Flegr J, Havlicek J, Kodym P, Maly M, Smahel Z. 2002. Increased risk of traffic accidents in subjects with latent toxoplasmosis: a retrospective case-control study. BMC Infect Dis 2:11.

13. Kocazeybek B, Oner YA, Turksoy R, Babur C, Cakan H, Sahip N, Unal A, Ozaslan A, Kilic S, Saribas S, Aslan M, Taylan A, Koc S, Dirican A, Uner HB, Oz V, Ertekin C, Kucukbasmaci O, Torun MM. 2009. Higher prevalence of toxoplasmosis in victims of traffic accidents suggest increased risk of traffic accident in Toxoplasma-infected inhabitants of Istanbul and its suburbs. Forensic Sci Int 187:103–8.

14. Flegr J, Klapilova K, Kankova S. 2014. Toxoplasmosis can be a sexually transmitted infection with serious clinical consequences. Not all routes of infection are created equal. Med Hypotheses 83:286–9.

15. Dunn D, Wallon M, Peyron F, Petersen E, Peckham C, Gilbert R. 1999. Mother-to-child transmission of toxoplasmosis: risk estimates for clinical counselling. Lancet 353:1829–33.

16. Dass SA, Vasudevan A, Dutta D, Soh LJ, Sapolsky RM, Vyas A. 2011. Protozoan parasite Toxoplasma gondii manipulates mate choice in rats by enhancing attractiveness of males. PLoS One 6:e27229.

17. Abdulai-Saiku S, Tong WH, Vyas A. 2017. Sexual Transmission of Cyst-Forming Coccidian Parasites with Complex Life Cycles. Current Sexual Health Reports 9:271–276.

18. Arantes TP, Lopes WD, Ferreira RM, Pieroni JS, Pinto VM, Sakamoto CA, Costa AJ. 2009. Toxoplasma gondii: Evidence for the transmission by semen in dogs. Exp Parasitol 123:190–4.

19. Dubey JP, Sharma SP. 1980. Prolonged excretion of Toxoplasma gondii in semen of goats. Am J Vet Res 41:794–5.

20. de Moraes EP, Batista AM, Faria EB, Freire RL, Freitas AC, Silva MA, Braga VA, Mota RA. 2010. Experimental infection by Toxoplasma gondii using contaminated semen containing different doses of tachyzoites in sheep. Vet Parasitol 170:318–22.

21. Hlaváčová J, Flegr J, Řežábek K, Calda P, Kaňková Š. 2021. Male-to-Female Presumed Transmission of Toxoplasmosis Between Sexual Partners. Am J Epidemiol 190:386–392.

22. Kaňková Š, Hlaváčová J, Flegr J. 2020. Oral sex: A new, and possibly the most dangerous, route of toxoplasmosis transmission. Med Hypotheses 141:109725.

23. Pena HFJ, Moroz LR, Sozigan RKB, Ajzenberg D, Carvalho FR, Mota CM, Mineo TWP, Marcili A. 2014. Isolation and Biological and Molecular Characterization of Toxoplasma gondii from Canine Cutaneous Toxoplasmosis in Brazil. Journal of Clinical Microbiology 52:4419–4420.

24. Odell AV, Tran F, Foderaro JE, Poupart S, Pathak R, Westwood NJ, Ward GE. 2015. Yeast three-hybrid screen identifies TgBRADIN/GRA24 as a negative regulator of Toxoplasma gondii bradyzoite differentiation. PLoS One 10:e0120331.

25. Flegr J, Prandota J, Sovičková M, Israili ZH. 2014. Toxoplasmosis – A Global Threat. Correlation of Latent Toxoplasmosis with Specific Disease Burden in a Set of 88 Countries. PLOS ONE 9:e90203.

26. Flegr J, Escudero DQ. 2016. Impaired health status and increased incidence of diseases in Toxoplasma-seropositive subjects - an explorative cross-sectional study. Parasitology 143:1974–1989.

27. Torgerson PR, Mastroiacovo P. 2013. The global burden of congenital toxoplasmosis: a systematic review. Bull World Health Organ 91:501–8.

28. Sarkar MD, Anuradha B, Sharma N, Roy RN. 2012. Seropositivity of toxoplasmosis in antenatal women with bad obstetric history in a tertiary-care hospital of Andhra Pradesh, India. J Health Popul Nutr 30:87–92.

29. Boyer KM, Holfels E, Roizen N, Swisher C, Mack D, Remington J, Withers S, Meier P, McLeod R. 2005. Risk factors for Toxoplasma gondii infection in mothers of infants with congenital toxoplasmosis: Implications for prenatal management and screening. Am J Obstet Gynecol 192:564–71.

30. Petersen E, Vesco G, Villari S, Buffolano W. 2010. What do we know about risk factors for infection in humans with Toxoplasma gondii and how can we prevent infections? Zoonoses Public Health 57:8–17.

31. Santana LF, Rossi GAM, Gaspar RC, Pinto VMR, de Oliveira GP, da Costa AJ. 2013. Evidence of sexual transmission of Toxoplasma gondii in goats. Small Ruminant Research 115:130–133.

32. Lopes WD, Rodriguez JD, Souza FA, dos Santos TR, dos Santos RS, Rosanese WM, Lopes WR, Sakamoto CA, da Costa AJ. 2013. Sexual transmission of Toxoplasma gondii in sheep. Vet Parasitol 195:47–56.

33. Dym M. 1994. Basement Membrane Regulation of Sertoli Cells*. Endocrine Reviews 15:102–115.

34. Siu MKY, Cheng CY. 2008. Extracellular Matrix and Its Role in Spermatogenesis. Advances in experimental medicine and biology 636:74–91.

35. Mruk DD, Cheng CY. 2015. The Mammalian Blood-Testis Barrier: Its Biology and Regulation. Endocrine Reviews 36:564–591.

36. Mital P, Hinton BT, Dufour JM. 2011. The Blood-Testis and Blood-Epididymis Barriers Are More than Just Their Tight Junctions. Biology of Reproduction 84:851–858.

37. Cheng CY, Mruk DD. 2010. A local autocrine axis in the testes that regulates spermatogenesis. Nat Rev Endocrinol 6:380–95.

38. Manavi K, Turner K, Scott GR, Stewart LH. 2005. Audit on the management of epididymo-orchitis by the Department of Urology in Edinburgh. Int J STD AIDS 16:386–7.

39. Trojian TH, Lishnak TS, Heiman D. 2009. Epididymitis and orchitis: an overview. Am Fam Physician 79:583–7.

40. Singhal S, Wagh DD, Kashikar S, Lonkar Y. 2011. A case of acute epididymo-orchitis due to Pseudomonas aeruginosa presenting as ARDS in an immunocompetent host. Asian Pacific Journal of Tropical Biomedicine 1:83–84.

41. Lim A, Kumar V, Hari Dass SA, Vyas A. 2013. Toxoplasma gondii infection enhances testicular steroidogenesis in rats. Mol Ecol 22:102–10.

42. Flegr J, Lindová J, Kodym P. 2008. Sex-dependent toxoplasmosis-associated differences in testosterone concentration in humans. Parasitology 135:427–31.

43. Hodkova H, Kolbekova P, Skallova A, Lindova J, Flegr J. 2007. Higher perceived dominance in Toxoplasma infected men--a new evidence for role of increased level of testosterone in toxoplasmosis-associated changes in human behavior. Neuro Endocrinol Lett 28:110–4.

44. Kumar V, Vasudevan A, Soh LJ, Le Min C, Vyas A, Zewail-Foote M, Guarraci FA. 2014. Sexual attractiveness in male rats is associated with greater concentration of major urinary proteins. Biol Reprod 91:150.

45. Tan D, Vyas A. 2016. Toxoplasma gondii infection and testosterone congruently increase tolerance of male rats for risk of reward forfeiture. Horm Behav 79:37–44.

